# A chromosome-level genome assembly of the classical model nematode *Panagrellus redivivus* for comparative studies of parasitism evolution

**DOI:** 10.64898/2026.10.22.761182

**Authors:** Kazuki Sato, Simo Sun, Tatsuya Yamashita, Akemi Yoshida, Ryoji Shinya, Paul W Sternberg, Taisei Kikuchi

## Abstract

*Panagrellus redivivus* is a free-living nematode that served as an early model system for nematode biology prior to *Caenorhabditis elegans*. As a member of Clade IV, a clade harboring major plant-parasitic (e.g., *Bursaphelenchus xylophilus*) and animal-parasitic (e.g., *Strongyloides* spp.) species, *P. redivivus* represents a crucial, phylogenetically proximate non-parasitic reference for investigating the evolution of parasitism. Here, we present a chromosome-level genome assembly of *P. redivivus* constructed using Oxford Nanopore long reads, Illumina short reads, and Hi-C scaffolding. The assembly is resolved into five chromosome-scale scaffolds, with completeness validated by Hi-C contact maps and the identification of a non-canonical telomeric repeat motif. The assembly demonstrates high gene-space completeness (97.1% BUSCO) and contains 17,839 predicted protein-coding genes. Comparative orthology analysis revealed extensive orthogroup sharing between *P. redivivus* and *Clade IV* parasites, enabling comparative evolutionary studies with reduced confounding effects from deep lineage divergence. This high-quality genome provides a key resource for investigating the genetic basis of distinctive biological traits, including ovoviviparity, as well as gene family dynamics and parasite evolution.

## Introduction

*Panagrellus redivivus* is a free-living nematode species that has long attracted interest in nematology and has been studied as an experimental model alongside *Caenorhabditis elegans* [1–3] (Fig. 1A). Comparative studies of *P. redivivus* and *C. elegans* have provided important insights into the evolution of nematode development, particularly through analyses of gonadal and postembryonic cell lineages [1, 2]. More recently, genomic and transcriptomic resources have further established *P. redivivus* as a useful comparative system for investigating the evolution of developmental and behavioural processes [4]. Studies comparing these two species have revealed both conserved and divergent features of cell lineages and behaviour [1, 2, 5]. In particular, *P. redivivus* differs from the oviparous *C. elegans* in its reproductive strategy, exhibiting ovoviviparity, in which embryos develop within the uterus and larvae are subsequently released from the female [6, 7].

**Fig. 1.**
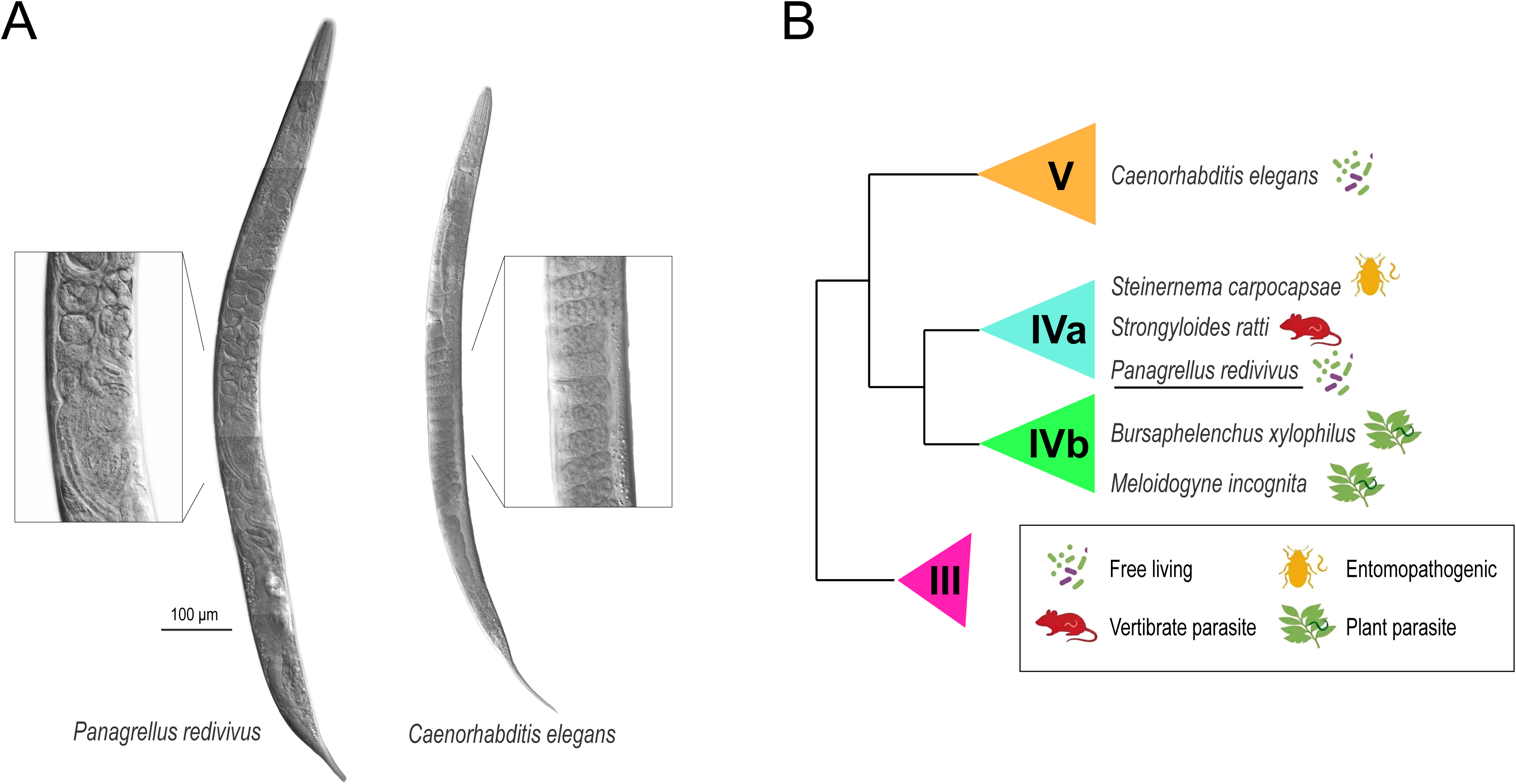
*Panagrellus redivivus* as a free-living comparative nematode model. (A) Light microscopy of an adult *P. redivivus* female and a *C. elegans* hermaphrodite. *P. redivivus* exhibit a slightly longer body length (∼1.5 mm) compared to *C. elegans (∼1.0 mm).* and exhibits ovoviviparity, in which eggs develop and hatch within the maternal uterus before the larvae are released. Magnified insets show developing embryos and larvae within the uterus of the adult *P. redivivus* female, highlighting this distinctive reproductive mode. In contrast, *C. elegans* is oviparous and releases eggs that subsequently develop and hatch outside the maternal body. Scale bar=100 µm. (B) Schematic species tree showing the phylogenetic relationships of nematode Clades III, IV, and V, adapted from Dorris et al [9]. Icons designate lifestyles: free-living, entomopathogenic, vertebrate parasitic, and plant parasitic. *P. redivivus* (underlined) is a free-living species situated within Clade IVa alongside parasitic relatives such as *S. ratti*.

While *C. elegans* belongs to Clade V of the nematode phylogeny, *P. redivivus* is a free-living member of the family Panagrolaimidae within Clade IV[8, 9] (Fig. 1B). Clade IV comprises a diverse assemblage of nematodes with free-living and parasitic lifestyles, including animal-parasitic lineages such as the genus *Strongyloides* and entomopathogenic nematodes of the genus *Steinernema* in Clade IVa, as well as plant-parasitic taxa such as *Bursaphelenchus xylophilus* in Clade IVb[8, 9]. This ecological and biological diversity makes Clade IV a valuable phylogenetic framework for investigating the evolutionary transitions between free-living and parasitic lifestyles in nematodes. In particular, the phylogenetic placement of the free-living *P. redivivus* provides a useful comparative system for examining the genetic and molecular changes associated with the evolution of parasitism. Phylogenetically informed comparisons between free-living and parasitic lineages can help distinguish conserved features from lineage-specific adaptations associated with host–parasite interactions and life-history strategies.

Despite its importance as a comparative model, *P. redivivus* remains relatively underexplored at the genomic level compared with extensively studied nematodes such as *C. elegans*. A draft genome and transcriptome of *P. redivivus* were previously generated, providing an important genomic resource for studies of nematode evolution and biology[4]. However, the limitations of this draft assembly have constrained investigations requiring a more complete representation of genome structure, including detailed analyses of chromosome organisation and evolutionary history. The earlier assembly was generated primarily from short-read sequencing data and was highly fragmented, limiting the resolution of genomic features and precluding robust chromosome-scale analyses.

Here, we present a high-quality chromosome-level genome assembly of *P. redivivus* generated using a hybrid sequencing strategy integrating Oxford Nanopore long reads, Illumina short reads for error correction, and Hi-C scaffolding. Similar hybrid approaches have proved highly effective in establishing chromosome-level reference genomes across diverse non-model invertebrates [10–12]. In *P. redivivus*, this approach substantially improves genome contiguity and chromosome-level resolution compared with the previously available draft genome. Notably, the chromosome-level assembly revealed non-canonical telomeric repeat sequences, providing new insights into chromosome-end organization in *P. redivivus*. This genome resource provides a robust foundation for investigating gene family evolution, chromosome evolution, and the genomic basis of nematode lifestyle transitions, particularly the evolution of parasitism within Clade IV.

## Methods

### Sample collection and DNA extraction

The *P. redivivus* strain PS2298/MT8872[1] used in this study was maintained in laboratory culture on standard nematode growth medium (NGM) seeded with *Escherichia coli* OP50 at 20°C. Nematodes were harvested from 14-day-old cultures, washed with M9 buffer, and stored at -80°C prior to DNA extraction. High-molecular-weight DNA was extracted from frozen nematode samples using lysis buffer containing 40 µg/ml proteinase K and 40 mM dithiothreitol. Following phenol-chloroform extraction, DNA was precipitated with isopropanol and resuspended in TE buffer. DNA quality and concentration were assessed using TapeStation (Agilent) and a Qubit 2.0 Fluorometer (Life Technologies).

### Genome sequencing and assembly

An initial genome assembly was generated using a hybrid sequencing strategy combining Oxford Nanopore long-read sequencing and Illumina short-read sequencing. High–molecular-weight genomic DNA was sequenced on an Oxford Nanopore MinION platform (Oxford Nanopore Technologies) using using R9-series flow cell, yielding approximately 6.1 Gb of long-read data with an N50 read length of 13.2 kb. In parallel, Illumina paired-end sequencing (PE; 2 × 300 bp) was performed on an Illumina platform. Combined with previously generated Illumina data, a total of 5.8 Gb of short-read data was obtained, providing approximately 90-fold coverage based on an estimated genome size of 65 Mb [4]. To achieve a chromosome-level assembly, Hi-C libraries were prepared using the Arima-HiC+ kit (Arima Genomics Inc.) to capture chromatin conformation and facilitate scaffolding. Hi-C libraries were sequenced as paired-end reads on an Illumina platform, generating approximately 4.8 million read pairs. De novo genome assembly was conducted using NextDenovo (v2.5.2) [13], followed by three rounds of base-error correction using Pilon (v1.24)[14] with Illumina short reads. Hi-C data were processed using Juicer (v1.5.7)[15] and the 3D-DNA pipelines (v180922)[16] resulting in a chromosome-level assembly. Assembly completeness was assessed using BUSCO (v5.8.2) [17] with the nematoda_odb12.2 lineage dataset.

K-mer frequency analysis was performed using KMC, a disk-based k-mer counting program designed for large sequencing datasets [18] and GenomeScope 2.0 [19]. The analysis was performed with a k-mer length of 21 and a diploid model.

### Genome annotation

Gene prediction was performed using the BRAKER2 pipeline (v2.1) [20], integrating RNA-seq data and protein homology information. RNA-seq data from mixed-stage nematodes [4] were used to guide splice site annotation. For protein homology support, a curated protein dataset comprising predicted proteins from Caenorhabditis elegans (WormBase WS280) and three Clade IV nematodes (*Bursaphelenchus xylophilus*, *Strongyloides ratti*, and the previous *P. redivivus* annotation) was provided to BRAKER2. Gene prediction was performed using the BRAKER2 pipeline (v2.1) [20], which integrates RNA-seq data and protein homology information to generate accurate gene models. RNA-seq data from mixed-stage nematodes [4] were used to guide gene annotation. For protein homology support, a curated protein dataset comprising predicted proteins from representative nematode genomes (∼10 species per clade across major nematode clades) retrieved from WormBase (release WS280) and WormBase ParaSite (release WBPS19; https://parasite.wormbase.org) was provided to BRAKER2. Functional annotation was conducted using EggNOG-mapper (v2.1.6) [21]. Repetitive elements were identified using RepeatModeler/RepeatMasker (v2.0.1/v4.1.8) [22] (https://www.repeatmasker.org/).

### Detection of telomeric repeat motifs from long-read sequences

Telomeric repeat motifs were detected from long-read sequencing data. Nanopore reads were error-corrected using Illumina short reads using FMLRC2 (v0.1.4)[23] to reduce sequencing errors prior to repeat detection and subsequently analysed using TeloSearchLR (v1.0.0)[24] with parameters -k 4 -K 20 and -n 6000.

### Identification of orthologous groups and species phylogeny

Gene family evolution and orthologous relationships were investigated using OrthoFinder (v2.5.5) [25]. Protein-coding gene sets from *P. redivivus* and three other nematode species—*C. elegans* (Clade V, free-living), *B. xylophilus* (Clade IV, plant-parasitic), and *Strongyloides ratti* (Clade IV, animal-parasitic)—were included in the analysis. Protein sequences were retrieved from WormBase ParaSite (WBPS19; https://parasite.wormbase.org). OrthoFinder was run with default parameters using the longest isoform per gene to identify orthogroups, single-copy orthologues, and gene family expansions or contractions. Orthogroup distributions were visualised using the R package UpSetR (1.4.0; R v4.4.1) [26] based on gene count outputs generated by OrthoFinder. To infer the phylogenetic relationships among nematode species, amino acid sequences from 4,166 single-copy orthologous genes were aligned separately using MAFFT (v7.526) [27] and trimmed using Gblocks (v1.0) [28]. Three alignments containing no retained amino acid sites after trimming were excluded, and the remaining 4,163 alignments were concatenated into a single alignment. The concatenated amino acid alignment was used to infer a partitioned maximum-likelihood phylogenetic tree using IQ-TREE (v2.4.0) [29]. The best-fit amino acid substitution model was selected using ModelFinder, and branch support was assessed using 1,000 ultrafast bootstrap replicates. Branch lengths were estimated as the number of amino acid substitutions per site.

### Nigon-based chromosomal analysis

Based on the OrthoFinder analysis, 5,375 genes were identified as one-to-one orthologous relationship between *C. elegans* and *P. redivivus*. Of these, 1,597 orthologs were assigned to seven Nigon elements (A–E, N, and X) [30]. Orthology data, genomic coordinates, and Nigon element assignments were integrated using the load_orthologs() function in the R package macrosyntR (https://github.com/SamiLhll/macrosyntR). Finally, the distribution of these Nigon-assigned genes along each *P. redivivus* chromosome was visualised using ggplot2 [31].

## Results and Discussion

### Chromosome-scale genome assembly

To characterize the *P. redivivus* genome prior to assembly, we analysed the k-mer frequency distribution of Nanopore sequencing reads using GenomeScope 2.0 with k = 21. The analysis estimated a total haploid genome size of 71.0 Mb and a heterozygosity rate of 2.05%, with a primary homozygous k-mer coverage peak at approximately 27.4× (Fig. 2A). The observed k-mer distribution was well fitted by the GenomeScope model, with an estimated sequencing error rate of 2.5%. We then generated a chromosome-level genome assembly of *P. redivivus* by integrating Oxford Nanopore long-read, Illumina short-read, and Hi-C sequencing data. The initial assembly, constructed from Nanopore long reads and polished with Illumina short reads, comprised 32 contigs with lengths ranging from 0.99 to 9.9 Mb. Taxonomic contamination screening confirmed high sequence purity, with the vast majority of sequence length assigned to Nematoda (Fig. S1). Subsequent Hi-C scaffolding successfully resolved these contigs into five major scaffolds corresponding to the expected five chromosomes of *P. redivivus* (Fig. 2B), yielding a total assembly size of 66.9 Mb (Table 1 and Fig. S2). This assembled size accounts for approximately 94.2% of the GenomeScope-estimated genome size (71.0 Mb), with the remaining ∼4.1 Mb difference (6.1%).

**Fig. 2.**
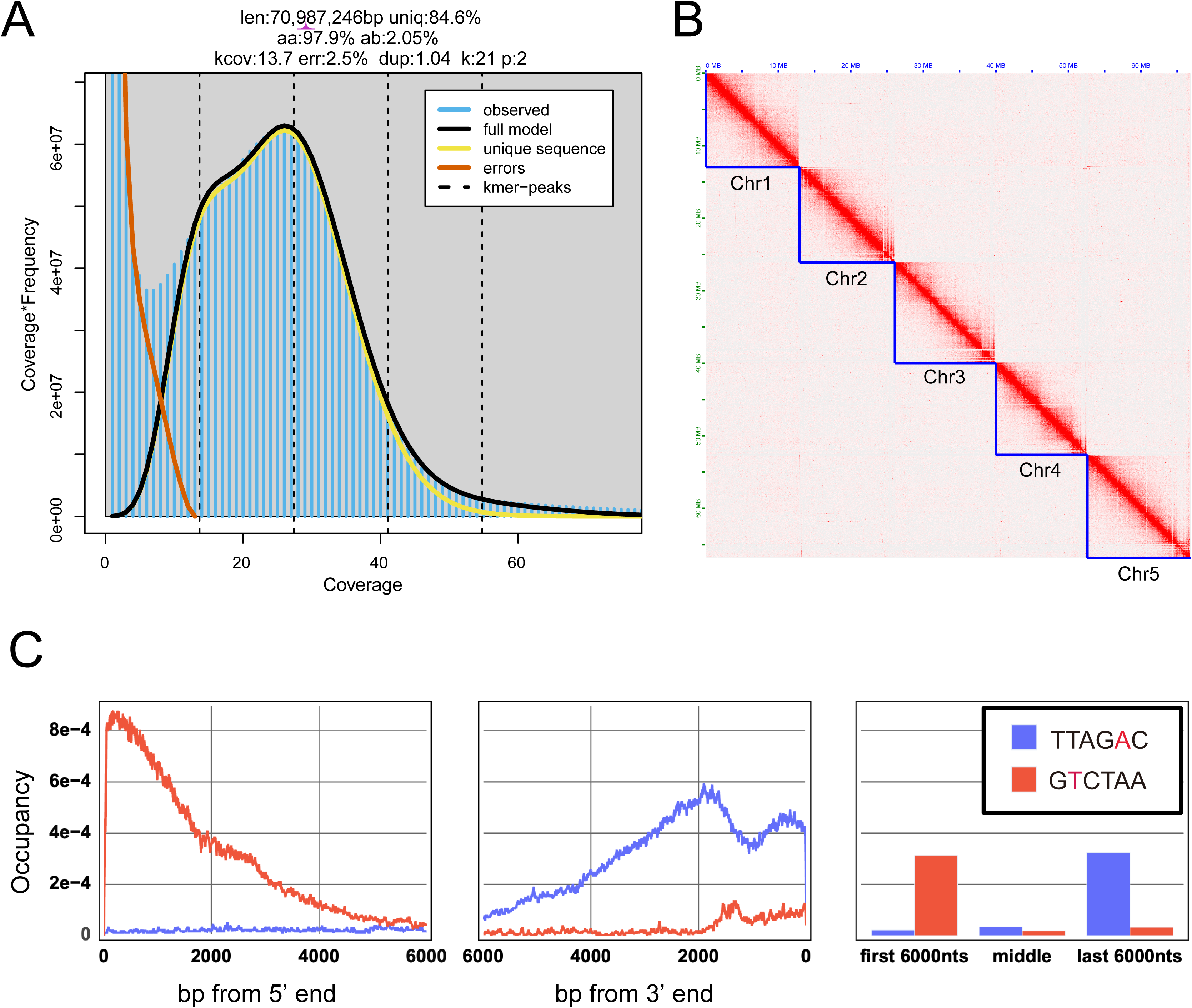
K-mer profile, chromosome-scale assembly, and terminal repeat composition of*Panagrellus redivivus*. (A) GenomeScope 2.0 analysis of the *P. redivivus* k-mer frequency distribution (k = 21). Observed k-mer frequencies (blue) are shown with the fitted model (black), unique-sequence component (yellow), sequencing-error component (orange), and k-mer peaks (dashed lines). The estimated genome size was 70.99 Mb, with 2.05% heterozygosity and a k-mer coverage of 13.7×. (B) Hi-C contact map of the chromosome-scale assembly, showing five distinct intrachromosomal interaction domains corresponding to chromosomes Chr1–Chr5. (C) Distribution of the canonical telomeric repeat TTAGAC (blue) and the non-canonical repeat GTCTAA (red) at chromosome termini. Repeat occupancy was quantified within the first and last 6 kb of each chromosome and compared with the middle regions.

**Table 1.** Statistics of the *P redivivus* genome assembly.

| Category | Metric | This study | Srinivasan *et al.*<br>(2013) |
| --- | --- | --- | --- |
| Genome assembly | Assembly size (Mb) | 66.9 | 64.4 |
|  | No. of chromosomes | 5 | - |
|  | GC content (%) | 44.1 | 44.3 |
|  | Repetitive sequences (%) | 11.59 | 7.01 |
|  | BUSCO genome completeness * | 96.0 (S: 94.1%, D: 1.9%) | 96.0 (S: 93.8%, D: 2.2%) |
| Gene prediction | No. of protein-coding genes | 17,839 | 24,249 |
|  | No. of protein-coding transcripts | 18,478 | 26,372 |
|  | Coding length (Mb)** | 23.82 | 26.85 |
|  | BUSCO protein completeness *** | 94.0% (S: 87.5%, D: 6.4%) | 95.5% (S: 83.3%, D: 12.2%) |

\* BUSCO completeness evaluated at the genome level using the nematoda\_odb12.2 dataset. S: single-copy BUSCOs; D: duplicated BUSCOs.
\*\* Total length of predicted coding sequences (CDS).
\*\*\* BUSCO completeness evaluated on predicted protein sequences (longest isoform per gene). S: single-copy BUSCOs; D: duplicated BUSCOs

### Quality and completeness of the assembly

BUSCO analysis indicated 96.0% genome completeness (94.1% single-copy and 1.9% duplicated BUSCOs; Table 1), demonstrating the high quality of the assembly. The assembled genome spanned 66.9 Mb, with 7.75 Mb (11.59%) annotated as repetitive sequences, a higher proportion than the 7.01% reported in the previous assembly by Srinivasan et al. [4] (Table 2). This increase likely reflects the improved resolution of complex repetitive regions enabled by long-read sequencing.

**Table 2.** Repeat elements of the *P. redivivus* genome.

|  | Length occupied (bp) | Percentage of sequence (%) |
| --- | --- | --- |
| Retroelements | 463,911 | 0.69 |
| SINEs | 19,707 | 0.03 |
| LINEs | 280,858 | 0.42 |
| LTR elements | 163,346 | 0.24 |
| DNA transposons | 520,457 | 0.78 |
| Rolling-circles | 41,545 | 0.06 |
| Small RNA | 95,571 | 0.14 |
| Satellites | 34,797 | 0.05 |
| Simple repeats | 245,977 | 0.37 |
| Low complexity | 58,006 | 0.09 |
| Unclassified | 6,293,251 | 9.41 |
| Total | 7,752,191 | 11.59 |

Gene prediction identified 17,839 protein-coding genes (18,478 transcripts), representing a substantial reduction from the 24,249 genes reported previously (Table 1). This decrease is primarily attributed to the resolution of fragmented gene models and false gene duplications that commonly artifact in short-read-based draft assemblies. Supporting this, the duplication rate in protein BUSCO analysis markedly dropped from 12.2% in the previous assembly to 6.4% in our study (Table 1), indicating a much cleaner, more consolidated gene set. Among the predicted genes, 7,789 (43.66%) were functionally annotated with Gene Ontology (GO) terms.

### Non-canonical telomeric repeat motifs in *P. redivivus*

We identified a non-canonical telomeric repeat motif (TRM), TTAGAC, in the chromosome-scale genome assembly of *P. redivivus.* A tandem repeat array exceeding 2.5 kb of the TTAGAC motif was detected at one end of chromosome 5, whereas the canonical nematode telomeric motif TTAGGC was not detected at any chromosomal termini. To independently validate this observation, we analysed Nanopore long-read sequencing data using TeloSearchLR[24]. This analysis revealed strong enrichment of TTAGAC repeats and their reverse complement (GTCTAA) at the termini of long reads, with motif frequency progressively decreasing toward internal read regions (Fig. 2C). These results provide independent long-read evidence supporting the localisation of TTAGAC repeats at chromosomal termini. Previous studies reported the emergence of TTAGAC repeats in a subset of Panagrolaimidae species, while other lineages, including the genus *Panagrolaimus,* retain the canonical TTAGGC motif [32]. Our chromosome-scale and long-read data provide direct evidence confirming the presence and terminal localisation of this non-canonical TRM in *P. redivivus*.

### Evolutionary suitability of *P. redivivus* as a comparative model: orthologue sharing and chromosome evolution

OrthoFinder analysis revealed extensive orthogroup (OG) sharing between *P. redivivus* and the Clade IV parasitic nematodes *B. xylophilus* (7,076 pairwise shared OGs) and *S. ratti* (6,587) (Fig. 3A). These shared orthologues provide a broad genomic basis for comparative analyses of gene family evolution between the free-living *P. redivivus* and parasitic Clade IV nematodes, including gene family expansion, contraction, and diversification. Comparable levels of OG sharing were also observed between *C. elegans* and the Clade IV parasitic species (*B. xylophilus*, 6,875; *S. ratti*, 6,706) (Fig. 3A). However, the extent of OG sharing may be influenced, at least in part, by annotation bias, because *C. elegans* proteins are frequently used as reference sequences in nematode gene prediction and annotation. To further resolve the evolutionary relationships underlying these shared gene repertoires, we inferred a species tree from 4,166 single-copy orthologues. This analysis recovered a strongly supported sister relationship between *P. redivivus* and *B. xylophilus* (local posterior probability = 1.0), with *S. ratti* as their next closest relative, whereas *C. elegans* was positioned outside the Clade IV lineages (Fig. 3B). Together, these results highlight the extensive conservation of gene repertoires between *P. redivivus* and Clade IV parasitic nematodes and establish *P. redivivus* as a valuable free-living comparative reference for investigating genomic changes associated with parasitism within this clade.

**Fig. 3.**
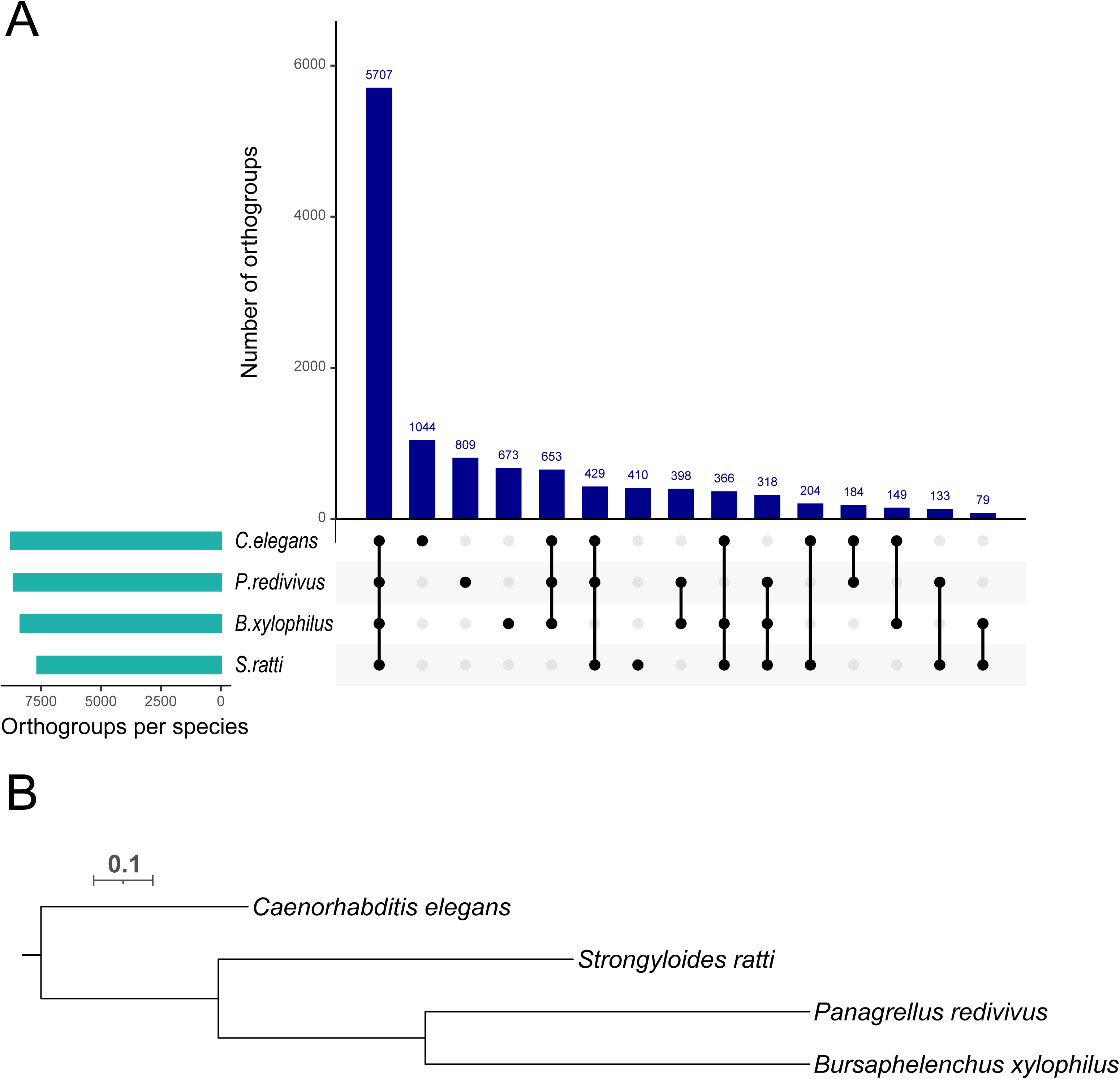
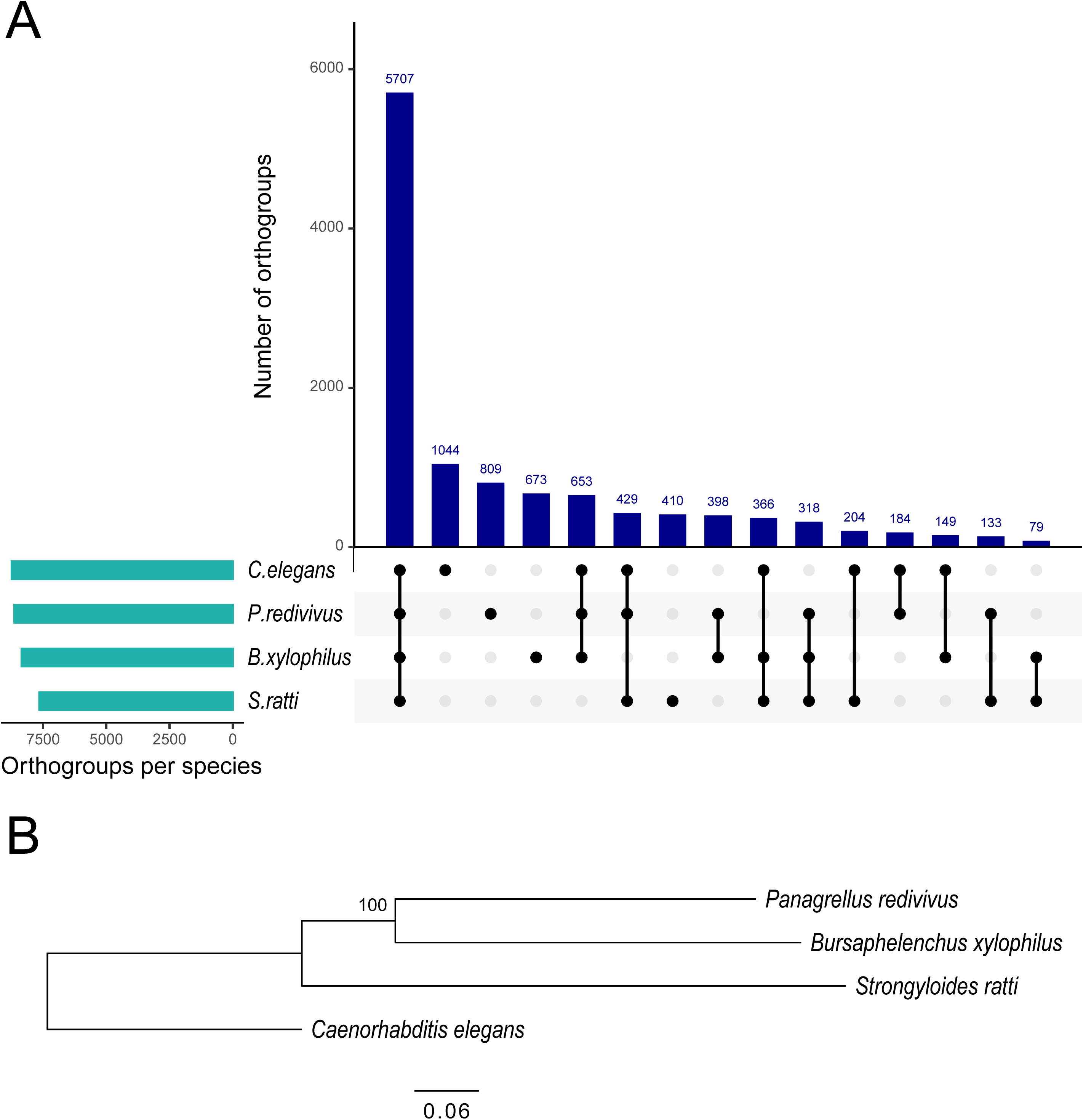
Orthogroup sharing and phylogenetic proximity of *Panagrellus redivivus* to Clade IV parasitic nematodes. (A) UpSet plot summarizing orthogroup distributions inferred by OrthoFinder among *C. elegans*, *P. redivivus*, *B. xylophilus*, and *S. ratti*. Dark blue bars indicate the numbers of orthogroups in each species-specific or shared orthogroup set, as defined by the black dots below. Light green bars indicate the total number of orthogroups identified in each species. (B) Phylogenetic tree inferred from a concatenated amino acid alignment of 4,163 single-copy genes. Branch lengths represent amino acid substitutions per site and indicate the relative evolutionary distances among species. *P. redivivus* and *B. xylophilus* form a close sister pair, with *S. ratti. C. elegans* shows progressively greater evolutionary distances from these species. Bootstrap support value is displayed at an internal node.

To investigate chromosome evolution in *P. redivivus*, we mapped 1,597 genes to the seven ancestral Nigon elements (nematode ancestral chromosome units). Nigon elements were distributed across all five chromosomes, with individual chromosomes containing contributions from multiple ancestral elements, revealing a mosaic organization of ancestral chromosomal segments (Fig. 4). Chromosomes 1, 2, 4, and 5 of *P. redivivus* predominantly correspond to the single Nigon elements A, B, E, and N, respectively. In contrast, chromosome 3 comprises a fusion of Nigon elements C, D, and X (Fig. 4). A similar Nigon organization has been reported in the six-chromosome genome of *B. xylophilus*, in which five chromosomes predominantly correspond to single Nigon elements, while chromosome 3 contains Nigon elements C and X [7, 30]. Given the close evolutionary relationship between *P. redivivus* and *B. xylophilus* (Fig. 3B), these patterns raise the possibility that a chromosome composed of Nigon elements C and X was present in their common ancestor. Under this scenario, the C+X chromosome would have been retained in the *B. xylophilus* lineage, whereas the *P. redivivus* lineage subsequently acquired an additional D-derived segment through chromosome fusion, giving rise to the present C+D+X chromosome. This proposed evolutionary history remains to be tested with broader taxon sampling and comparative chromosome-level analyses.

**Fig. 4.**
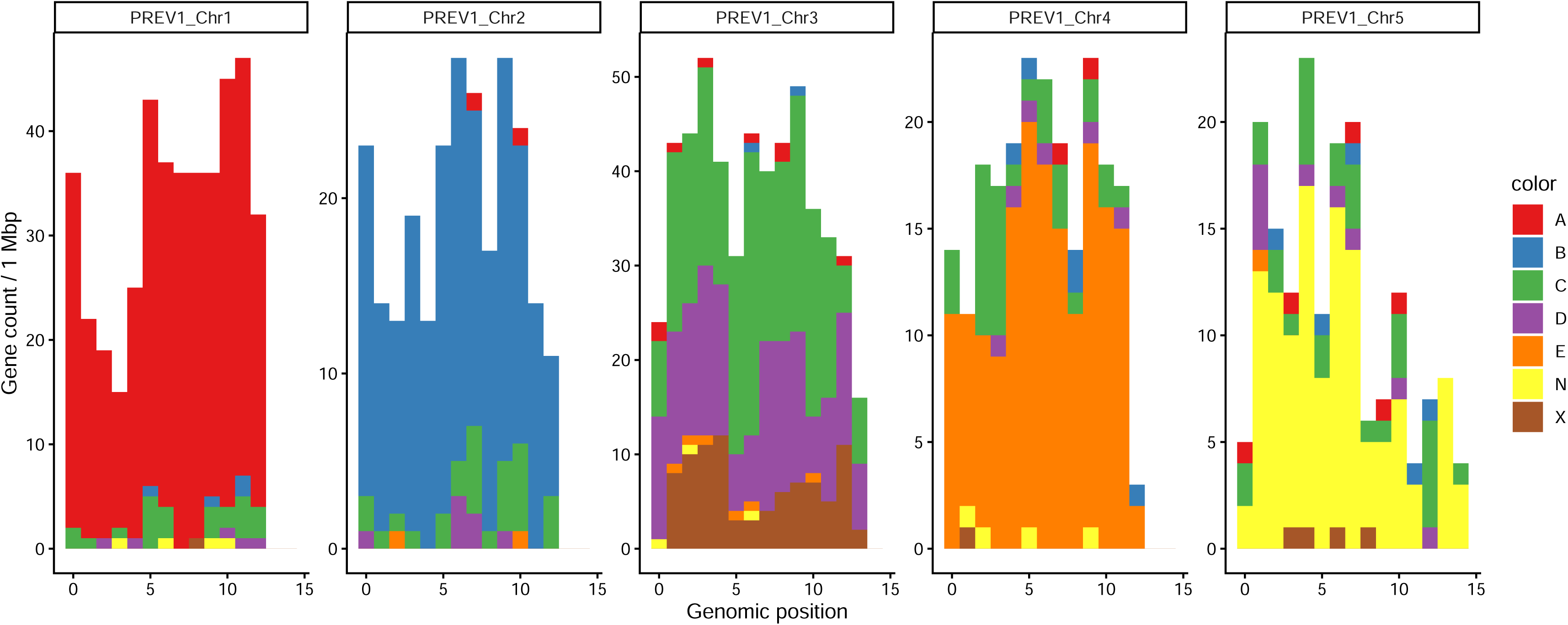
Nigon element composition of *Panagrellus redivivus* chromosomes. The distribution of 1,597 genes assigned to the seven ancestral Nigon elements across the five *P. redivivus* chromosomes. Gene counts are shown in 1-Mbp genomic windows, with each gene colored according to its assigned Nigon element. The mosaic distribution of Nigon elements across chromosomes indicates extensive reshuffling of ancestral chromosomal segments in the *P. redivivus* lineage.

### Availability of a chromosome-scale assembly of *P. redivivus*

The chromosome-scale genome assembly of *P. redivivus* presented here provides a high-quality resource for comparative studies of genome and chromosome evolution. Its position within Clade IV makes *P. redivivus* a useful free-living reference for comparisons with parasitic nematodes. The chromosome-scale resolution also enables detailed analyses of chromosome evolution, including changes in Nigon element organization. Together with the extensive resources available for *C. elegans*, this genome provides a complementary model for comparative and functional studies of nematode biology.

## Data availability

All sequencing reads are accessible through the International Nucleotide Sequence Database Collaboration (INSDC) repositories, including DRA, SRA and ENA, under accession numbers DRR1089823 / SRR25684730 (Illumina), DRR1089824 (Oxford Nanopore), and DRR546971 (Hi-C reads). The assembled genome and associated annotations are available in INSDC databases under BioProject accession PRJEB124666. The RNA-seq data used for gene annotation are accessible in the SRA under accession number SRR19570654.

## Declarations

## Ethics approval and consent to participate

The authors declare that ethical approval was not required for this type of research.

## Acknowledgments

We extend our gratitude to the members of the Parasite Systems Biology Lab in The University of Tokyo for their useful advice and continuous support.

## Author contributions

Kazuki Sato (Investigation, Formal analysis, Writing—original draft), Simo Sun (Investigation, Formal analysis), Tatsuya Yamashita (Resources, Visualization), Akemi Yoshida (Investigation, Resources), Ryoji Shinya (Resources, Conceptualization), Paul W Sternberg (Resources, Conceptualization), and Taisei Kikuchi (Conceptualization, Funding acquisition, Project administration, Supervision, Writing—review & editing)

