## Supplementary material for "A chromosome-level genome assembly of the classical model nematode *Panagrellus redivivus* for comparative studies of parasitism evolution": Fig. S1

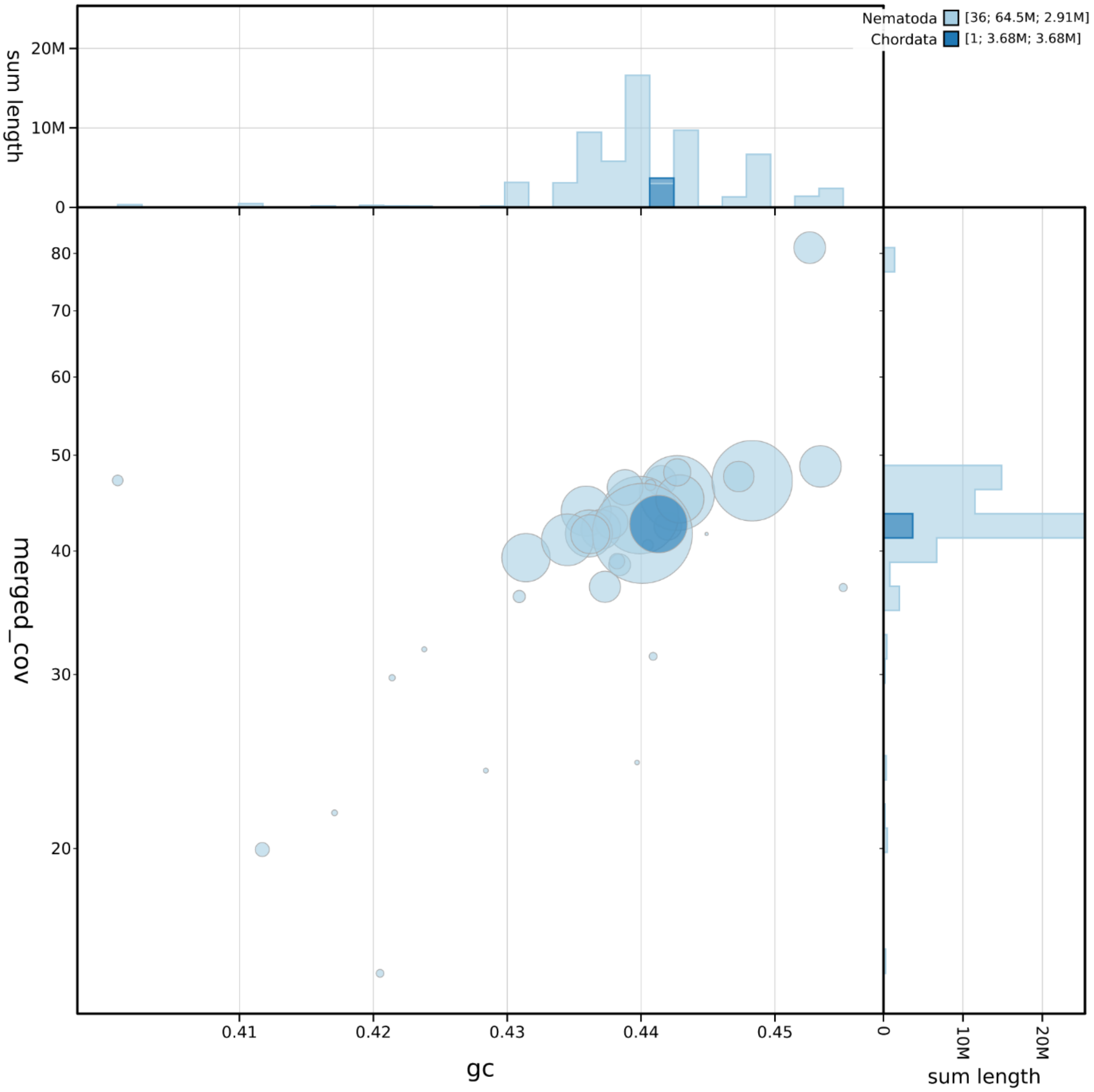


Supplementary Figure S1. Taxonomic contamination screening of the *Panagrellus redivivus* genome initial assembly generated using Nanopore reads. BlobToolKit blob plot displaying the GC content (x-axis) versus sequencing coverage (y-axis) based on Illumina reads for the genome scaffolds. Each circle represents a scaffold, scaled proportionally to its length and coloured according to phylum-level taxonomic assignment based on BLAST searches. All contigs belong to Nematoda (light blue; 36 contigs, total length 64.5 Mb) except one that is assigned to Chordata (dark blue; 3.68 Mb). Marginal histograms on the top and right depict the distribution of total assembly length as a function of GC content and read coverage, respectively.

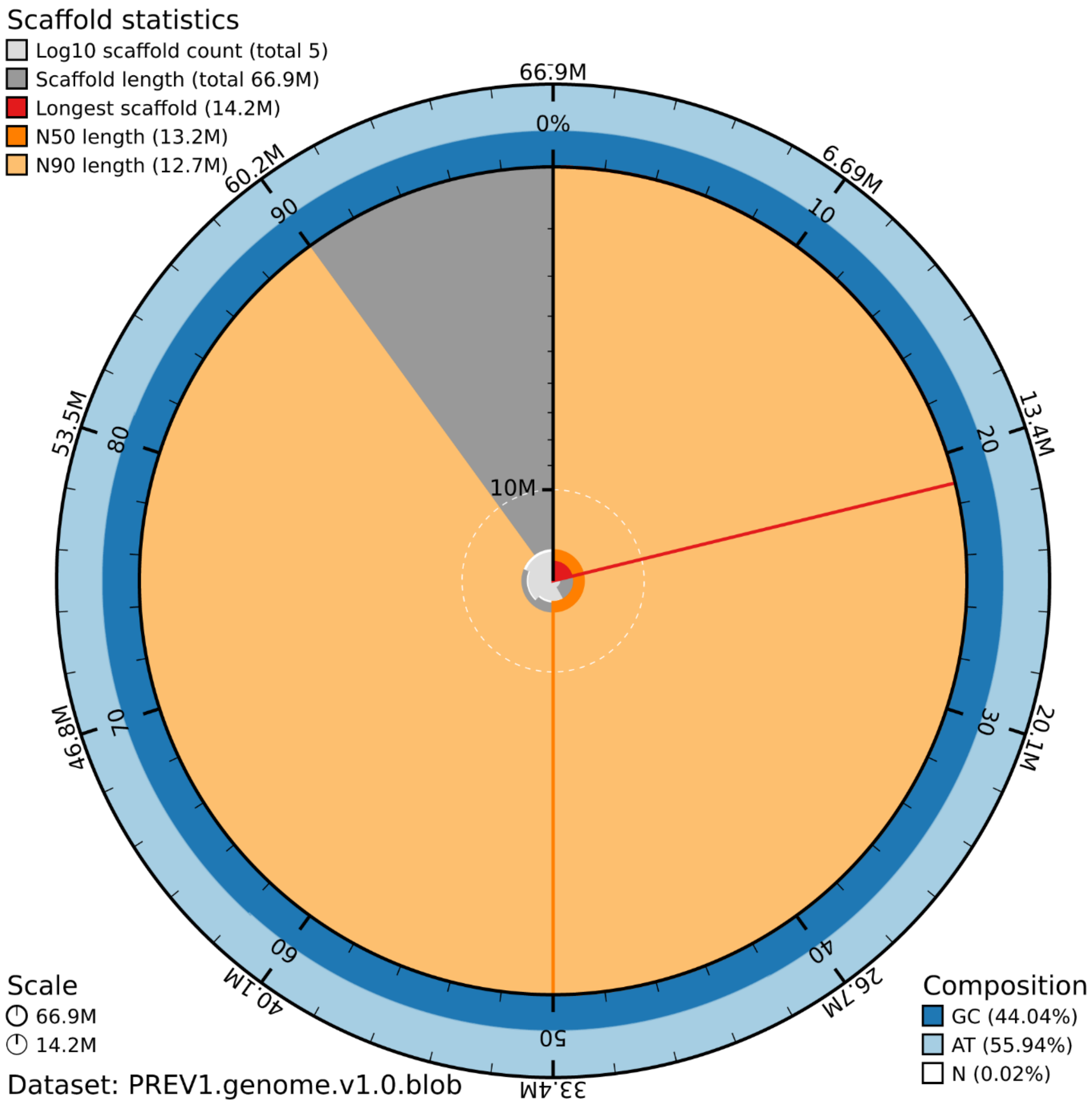


Supplementary Figure S2. Scaffold statistics and nucleotide composition of the *Panagrellus redivivus* genome assembly. BlobToolKit snail plot summarising assembly metrics for the *Panagrellus redivivus* genome (dataset: PREV1.genome.v1.0). The plot illustrates a highly contiguous assembly comprising 5 scaffolds with a total length of 66.9 Mb. The longest scaffold measures 14.2 Mb (indicated by the red line/arc), with an N50 length of 13.2 Mb (dark orange) and an N90 length of 12.7 Mb (light orange). The outer concentric rings represent base composition: 44.04% GC (dark blue), 55.94% AT (light blue), and 0.02% ambiguous bases (Ns; white).
